# Catecholaminergic axons in the neocortex of adult mice regrow following brain injury

**DOI:** 10.1101/737072

**Authors:** Sarah E. Dougherty, Tymoteusz J. Kajstura, Yunju Jin, Michelle H. Chan-Cortés, Akhil Kota, David J. Linden

**Affiliations:** The Solomon H. Snyder Department of Neuroscience, Johns Hopkins University School of Medicine. Baltimore, MD. USA; Department of Neurobiology and Anatomy, University of Utah, School of Medicine. Salt Lake City, UT, USA

**Keywords:** stab injury, controlled cortical impact, tyrosine hydroxylase, dopamine, acetylcholine, norepinephrine, regeneration

## Abstract

Serotonin axons in the adult rodent brain can regrow and recover their function following several forms of injury including controlled cortical impact (CCI), a neocortical stab wound, or systemic amphetamine toxicity. To assess whether this capacity for regrowth is unique to serotonergic fibers, we used CCI and stab injury models to assess whether fibers from other neuromodulatory systems can also regrow following injury. Using tyrosine-hydoxylase (TH) immunohistochemistry we measured the density of catecholaminergic axons before and at various time points after injury. One week after CCI injury we observed a pronounced loss, across cortical layers, of TH+ axons posterior to the site of injury. One month after CCI injury the same was true of TH+ axons both anterior and posterior to the site of injury. This loss was followed by significant recovery of TH+ fiber density across cortical layers, both anterior and posterior to the site of injury, measured three months after injury. TH+ axon loss and recovery over weeks to months was also observed throughout cortical layers using the stab injury model. Double label immunohistochemistry revealed that nearly all TH+ axons in neocortical layer 1/2 are also dopamine-beta-hyroxylase+ (DBH+; presumed norepinephrine), while TH+ axons in layer 5 are a mixture of DBH+ and dopamine transporter+ types. This suggests that noradrenergic axons can regrow following CCI or stab injury in the adult mouse neocortex and leaves open the question of whether dopaminergic axons can do the same.

**Highlights:**

- We measured catecholaminergic axon density using tyrosine hydroxylase immunohistochemistry following two forms of brain injury.
- Both controlled cortical impact and cortical stab injuries caused extensive damage to catecholaminergic axons in the neocortex of adult mice.
- Following both types of injury, axon density slowly returned to control values over many weeks, including, in the case of stab injury, regrowth across the stab rift.
- Together with previous results showing serotonin axon regrowth, these findings suggest that monaminergic axons have an unusual capacity for regrowth following injury in the adult mammalian brain

## Introduction

A recent review article opened with a statement that has become broadly accepted in the field: “Central nervous system (CNS) axons do not spontaneously regenerate after injury in adult mammals.” (Heubener and Strittmatter, 2009). While this may be true for most neurons in the mammalian CNS, it does not hold for axons of neurons using the monoamine neurotransmitter serotonin. Serotonin axons broadly innervate the brain and spinal cord. In damaged rodent spinal cord, repaired with grafts and treated with chondroitinase, immunohistochemical studies have shown that some serotonin axons originating from the caudal raphe nuclei have crossed the graft and contributed to the restoration of locomotion, as well as control of the bladder and diaphragm (Zhou et al., 1995; Lee et al., 2010; Alilain et al., 2011; Lee et al., 2013; Yoo et al., 2013; Kanno et al., 2014, see Perrin and Noristani, 2019, for review). Similarly, histological studies have shown that serotonin axons can regrow following several types of rodent brain injury including a thermal lesion (Hawthorne, et al. 2011), a stab injury (Jin et al., 2016) or a controlled cortical impact (CCI) delivered to the neocortex (Kajstura et al., 2018). Using repeated *in vivo* two-photon imaging of adult serotonin transporter-EGFP BAC transgenic mice (Gong et al., 2003), we found that, in the weeks following neocortical stab injury, a large fraction (>80 %) of the severed ends of serotonin axons display new growth and in many cases this regrowth is sufficient to completely cross the glial scar-laden stab rift (Jin et al., 2016).

Histological investigations have also shown that serotonin axons regrow throughout the brain following their destruction with repeated high doses of para-chloro-amphetamine (PCA; O’Hearn et al., 1988; Molliver et al., 1990; Mamounas et al., 2000; Jin et al., 2016), a manipulation that fails to produce a glial scar (Wilson and Molliver, 1994). This indicates that formation of a glial scar is not necessary for serotonin axon regrowth in the brain. *In vivo* imaging showed that serotonin axons underwent widespread long-distance retrograde degeneration following PCA treatment and that the subsequent slow recovery of axon density in the neocortex was dominated by long-distance regrowth with little contribution from local sprouting. Unlike regrowing axons in the peripheral nervous system (Scheib and Hoke, 2013), new serotonin axons did not follow the pathways left by degenerated axons. Following PCA lesion, the regrown serotonin axons entering the field of view largely recapitulated the varicosity density, tortuosity and layer-specific distribution of pre-lesion axons. These regrown axons also had the capacity to release serotonin as measured by fast-scanning voltammetry. Thus, serotonin axons in the brain have an unusual capacity to regrow. Furthermore, this regrowth is at least somewhat functional and, in time, comes to approximates the pre-lesion state (Jin et al., 2016).

The serotonin system has been broadly implicated in the regulation of affect, aggression, sexual behavior, thermoregulation, sleep and cognitive function (Muller and Jacobs, 2010). Importantly, the effects of serotonin on neurons are modulatory in nature and act on the time scale of hundreds of milliseconds to tens of seconds. The serotonin axons that innervate the neocortex are thin and unmyelinated and they originate in the dorsal and median raphe nuclei. They form a C-shape when observed in the sagittal plane of the mouse brain: they pass through the lateral hypothalamus and the basal forebrain, running in the medial forebrain bundle. The fibers then turn dorsally, elaborating branches that form projections to the frontal cortex, and then turn posteriorly to innervate parietal, temporal and occipital cortices in turn (Figure 1A; Tork, 1990; Vitalis et al., 2013).

**Figure 1.**
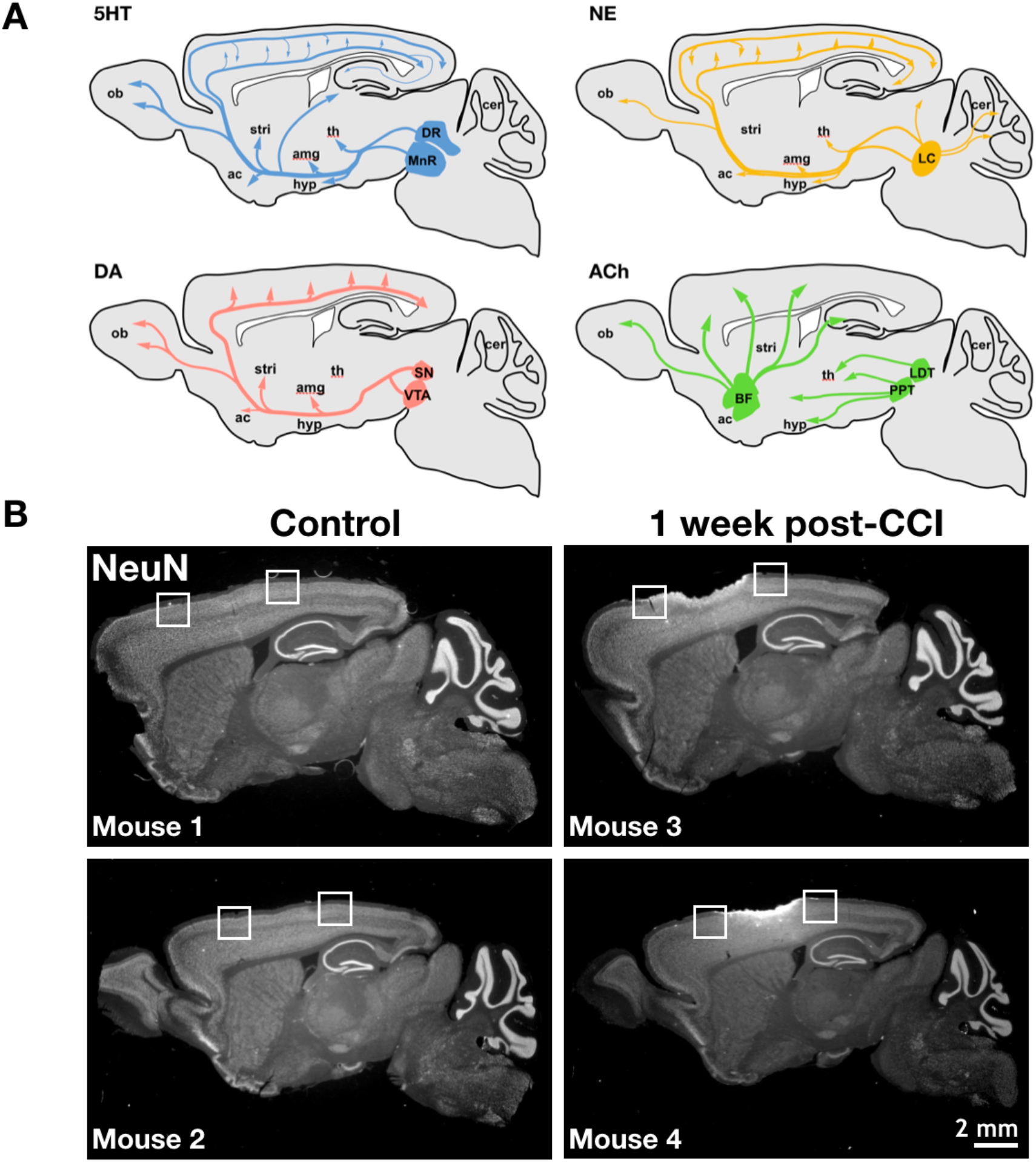
Neuromodulatory axons innervate the adult mouse forebrain, including the region injured by our CCI protocol. **A.** A schematic diagram showing the innervation of the forebrain by several neuromodulatory systems in a sagittal slice of adult mouse brain. Axons originating in the rostral serotonergic (5HT) nuclei, including the median raphe (MnR) and dorsal raphe (DR), project throughout the forebrain innervating regions including the neocortex, nucleus accumbens (ac), amygdala (amg), hypothalamus (hyp), olfactory bulb (ob), striatum (stri), and thalamus (th). Axons of the dopaminergic (DA) system originate in the ventral segmental area (VTA) and the substantia nigra (SN) and innervate the forebrain, including the neocortex, stri, and amg. The noradrenergic axons (NE) originate from neurons in the locus coeruleus (LC) and similarly project throughout the forebrain. The 5HT, NE and DA systems have a roughly similar trajectory: first coursing through the basal forebrain, ascending and then running into the cortex from anterior to posterior. Axons of the cholinergic (ACh) system that innervate the neocortex originate in a complex of nuclei in the basal forebrain (BF). They infiltrate the neocortical region through several ascending pathways. The pedunculopontine tegmental nucleus (PPT) and laterodorsal tegmental nucleus (LDT) predominately provide cholinergic innervation to the subcortical regions, midbrain and the BF. **B.** Low magnification exemplar images of sagittal sections, stained with the neuronal nuclear marker NeuN, reveal the extent of injury evoked by CCI in two exemplar mice per condition. One week after surgery the damage inflicted by controlled cortical impact is apparent. The control group mice show superficially intact neocortical regions. The white boxes show the regions for analysis of tissue flanking the injury site.

Is the ability of serotonin neurons in the adult brain to regrow their axons following injury shared by other types of neuron? If so, the axons of the other monamine neuromodulators such as the catecholamines, norepinephrine and dopamine, are good candidates. They also originate within nuclei of the midbrain/brainstem region, are unmyelinated (Beaudet and Descarries, 1978; Yeomans, 1989) and run in a roughly similar course to innervate the forebrain: through the medial forebrain bundle prior to ascending and then passing through the neocortex in the anterior-to-posterior direction (Figure 1A; Papadopoulos and Parnavelas, 1991; Nakamura et al., 2000; Berridge et al., 2003; Shnitko and Robinson, 2004; Sara, 2009; Juarez and Han, 2016).

Immunohistochemical experiments with antibodies directed against the catecholamine marker tyrosine hydroxylase (TH) show that catecholaminergic axons in the adult mammalian brain can regrow following a chemical lesion. Recovery of large-diameter TH+ fiber density, suggestive of regrowth, has been described after lesioning the mesostriatal dopamine system with MPTP (1-methyl-4-phenyl-1,2,3,6-tetrahydropyridine; Bezard et al., 2000; Song and Haber, 2000), 6-hydroxydopamine (Blanchard et al., 1996; Stanic et al., 2003) or methamphetamine (Granado et al., 2018). Similarly, following treatment of adult rats with the selective norepinephrine neurotoxin DSP-4, (N-(2-chloroethyl)-N-ethyl-2-bromobenzylamine), noradrenergic axon density of the neocortex, hippocampus and thalamus was initially depleted, but was substantially recovered 12 month later, as indicated by dopamine-beta-hydroxylase (DBH) immunohistochemistry (Fritchy and Grzanna, 1992). These findings suggest catecholaminergic fibers may share the regrowth properties of serotonin axons following chemical lesions. Whether there is catecholaminegic fiber regrowth following physical injury remains unknown.

Here, to test the hypothesis that catecholamine axons can regrow in the neocortex following physical brain injury, we have subjected adult mice to CCI or neocortical stab injuries, using the protocols we previously employed in studies of serotonin axon regeneration (Jin et al., 2016; Kajstura et al., 2018). We have performed double immunohistochemistry for TH and NeuN, a marker of neuronal cell bodies, at various time points after injury to assess the damage and potential regrowth of catecholamine axons.

## Methods

### Mice

All animal procedures were approved by the Institutional Animal Care and Use Committee at Johns Hopkins Medical Institute. For CCI experiments, surgeries were performed on female C57BL/6 mice (Charles River). For cortical stab injury experiments, we used male and female Slc6a4-EGFP BAC transgenic mice made by Dr. Charles Gerfen (NIH) as part of the GENSAT consortium (line RP23-39F11, BAC BX86, RRID: MMRRC_030692-UCD; Gong et al., 2003). For labelled norepinephrine neurons, BAC transgenic mice in which cre recombinase was driven by the promoter of the dopamine-β-hydroxylase gene (GENSAT, STOCK Tg(Dbh-cre)KH212Gsat/Mmucd, Stock #032081-UCD, RRID: MMRRC_032081-UCD; Gong et al., 2007) were crossed with the mTmG reporter line (Jackson Laboratory, Stock #007576, Gt(ROSA)26Sortm4(ACTB-tdTomato,-EGFP)Luo/J, RRID: MGI:3722405; Muzumdar et al., 2007). All mice were group housed at up to 5 mice per cage on a 12 hour light/dark cycle. Mice were given food and water *ad libitum*.

### Controlled Cortical Impact

CCI injury was performed as previously described (Kajstura et al., 2018). Briefly, mice were anesthetized with 3% isoflurane and placed in a stereotaxic device (Stoelting) with continued isoflurane delivery. The skin was retracted and fascia removed to expose the skull. A round craniotomy was made with an electric drill using a 2.7mm trephine drill bit (Fine Science Tools). The skull was removed and an electromagnetic impactor with a 1.5 mm tip (Leica Benchmark Stereotaxic Impactor model #39463920) was placed over the craniotomy site perpendicular to the brain surface in contact with the intact dura. The impactor was then retracted and then activated to impact the brain with a velocity of 4m/s, dwell time of 100 ms, and depth of 0.75 mm from the dural surface. Following impact injury, the skin was closed above the craniotomy using 9 mm wound clips (Fine Science Tools). Mice were given subcutaneous injections of Baytril (2.5 mg/kg) and Buprenex (0.01 mg/kg) and were monitored before being returned to group housing. Control mice received the same treatment (anesthesia, incision, staples, drug treatment) excluding the craniotomy and the impactor injury.

### Stab Injury

The stab injury was performed as previously described (Jin et al., 2016). Briefly, mice were anesthetized with 3% isoflurane and placed in a stereotaxic device (Stoelting) with continued isoflurane delivery. A small dose of dexamethasone (0.04 mL at 2 mg/mL) was administered by subcutaneous injection prior to surgery to minimize potential swelling at the surgical site, and a subcutaneous injection of buprenorphine (0.3 mg/mL) was also given before surgery to alleviate pain. A 2 by 2 mm craniotomy was made overlying the right somatosensory cortex. With the pial surface exposed, a miniature blade (Surgistar, USM 6700) was placed on the dura with a stereotaxic device at a location that was roughly centered in the window but which was chosen to avoid major surface blood vessels. The blade was then slowly advanced into the brain to the depth of 2 mm. The blade was immediately retracted and saline-soaked surgifoam (Johnson & Johnson) was applied to stop bleeding, if necessary. A square cover glass was placed over the craniotomy and sealed with Metabond dental cement (Parkell). Mice were given a subcutaneous injection of Baytril (2.5 mg/kg), monitored for distress and then returned to group housing.

### Tissue Preparation and Immunohistochemistry

Mice were anesthetized with ketamine (100 mg/kg) and xylazine (10 mg/kg). They were then sacrificed by intracardial perfusion with ice-cold 4% paraformaldehyde (PFA) in phosphate buffered saline (PBS). Brains were then removed, incubated in 4% PFA overnight at 4°C and then cryopreserved in graded sucrose steps (15% and 30% for 24 hours each) at 4°C. Whole brains were sectioned on a sliding microtome (Leica) at 40 μm thickness and stored in PBS with Na-Azide (Fisher) at 4°C until use. Brain sections were washed in 0.3% TritonX100 PBS and then blocked for 2 hours at room temperature in 5% normal goat serum. Sections were then incubated overnight at 4°C in primary antibodies. Double-labeling was performed with each marker of a neuromodulator and NeuN (to identify cortical layers). The primary antibodies used were rabbit anti-TH (Millipore #AB152, RRID: AB_390204), rabbit anti-VAChT (Synaptic Systems #139103, RRID: AB_887864), rat anti-DAT (Millipore #MAB369, RRID:AB_2190413), chicken anti-GFP (Aves Labs #GFP-1010, RRID: AB_2307313) and mouse anti-NeuN (Millipore #ABN60; RRID: AB_2298767). Primary antibody dilutions were all 1:1000 with the exception of the anti-GFP primary which was used at a dilution of 1:6000. Sections were then washed in .3% TritonX100 PBS followed by overnight incubation in secondary antibodies at 4°C. Secondary antibodies used were Alexa Fluor 488-labeled goat anti-chicken (Jackson ImmunoResearch Laboratories #103-007-008, RRID: AB_2632418), Alexa Fluor 594-labeled goat anti-mouse (Jackson ImmunoResearch Laboratories #115-586-146, RRID: AB_2338899), Alexa Fluor 647-labeled goat anti-rat (Jackson ImmunoResearch Laboratories #112-606-062; RRID: AB_2338407), Alexa Fluor 647-labeled goat anti-rabbit (Jackson ImmunoResearch Laboratories #111-607-008; RRID: AB_2632470), and Alexa Fluor 488-labeled goat antirabbit (Jackson ImmunoResearch Laboratories #111-547-003; RRID: AB_2338058). Secondary antibody dilutions were all 1:500. Sections were washed, mounted on slides, and coverslipped using Prolong Antifade Diamond mounting media. Full section thickness z-stack images were acquired using a laser scanning confocal microscope (Zeiss LSM 880 AxioExaminer.Z1) with a 1 μm step size.

### Image Analysis

Raw confocal images were obtained as czi files using Zeiss Zen Black software, exported as channel merged z-stack tifs and analyzed using custom code written using the Anaconda Python distribution (Continuum Analytics). Full z-stack maximum projection images were used to set a detection threshold for pixels positive for either TH or VAChT signal. NeuN staining was used as a cortical layer marker and an indicator of tissue quality. Using an iterative threshold detection algorithm (Kajstura et al., 2018), the intensity of all positive pixels were set to 255 (the highest value for an 8-bit image) and background pixels were set to 0. The fiber density quantification was expressed as the percent supra threshold pixels within image area.

We performed axon tracing on volume-matched maximum-projected image stacks using Imaris (Bitplane; Figure 8). TH+ axons were exhaustively traced within a 50 μm wide designated-rift area. Total axon length within stab rift area and the number of mid-rift axon crossings were measured. For colocalization studies, TH+ axon segments were exhaustively traced within a 75 x 75 μm^2^ region of interest using only the TH channel of the z-stack maximum projection image. These traces were then superimposed onto the corresponding DBH-GFP channel ROI and each segment was manually scored as either positive or negative for GFP. One-tailed student’s t-tests were used to evaluate significance in layer-specific single time points. ANOVA tests were performed for layer-specific comparisons over time and any comparison across three or more groups.

## Results

### TH+ axons in the neocortex were damaged by CCI

To test the hypothesis that catecholaminergic axons are damaged and then regrow following injury, we subjected adult mice to CCI delivered to the anterior somatosensory cortex or a control condition in which the incision was performed but without the craniotomy or the impact. Mice were sacrificed one week, one month, or three month after these treatments. Their brains were then fixed, sliced in the sagittal plane and processed for double-label immunohistochemistry using antibodies directed against TH as a marker of catecholaminergic axons and NeuN to reveal neuronal nuclei and thereby serve as a marker for cortical layer boundaries.

Confocal 35 μm thick stacks were acquired using a 40x objective in layers 1, 2/3, and 5 of the motor cortex, anterior to the injury, and the posterior somatosensory cortex, posterior to the injury. We purposely disregarded the portion of the cortex that was directly underneath the impactor and hence was crushed and unsuitable for analysis (Figure 1B).

In both CCI and control tissue, distinct varicosity-bearing TH+ axons were seen in all layers of the cortex, corresponding to regions anterior and posterior to the craniotomy site (Figure 2). The axons tend to run in the anterior-posterior direction, with this orientation being strongest in layer 1. These images are consistent with previous reports using TH immunohistochemistry on tissue from rats (Kritzer, 1998) humans, chimpanzees, and monkeys (Raghanti et al., 2008).

**Figure 2.**
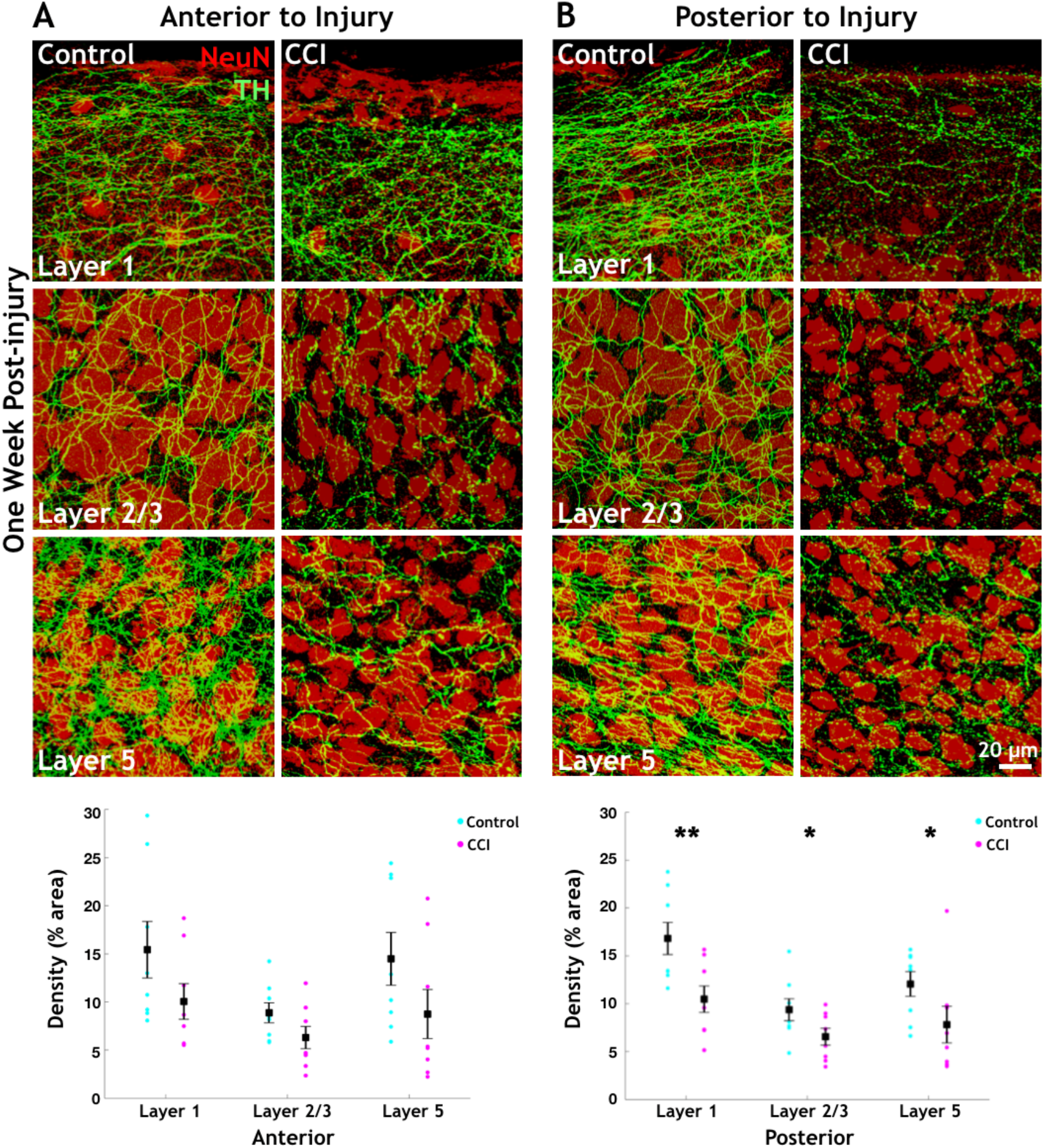
One week after CCI, the density of TH+ axons in the neocortex is reduced in the region posterior to the injury site as compared with controls. The exemplar images show maximally projected z-stacks with thresholded TH staining to indicate catecholaminergic axons and NeuN staining for cell bodies to clarify the boundaries of the cortical layers. **A.** Anterior to CCI injury, loss of TH+ axons is not statically significant in any cortical layer examined as compared to control tissue. Percent pixels positive for TH signal were calculated within the image area. Each circular plot point represents one animal, squares are the average within treatment group. Error bars are SEM. **B.** Posterior to the injury, significant TH+ axon density reductions are seen in cortical layer 1, layer 2/3, and layer 5. n= 8 animals/group. One-tailed Student’s t-test. * indicates p < 0.05. ** indicates p < 0.01. These and all images that follow are sagittal sections.

One week after injury, posterior to the injury site, statistically significant reductions in TH+ axon density were observed in all cortical layers examined when compared to controls. TH+ axon density in CCI injured tissue was reduced in layer 1 from 16.82% ± 1.67 to 10.48% ± 1.38 (1-tailed t-test, p = 0.0057), in layer 2/3 from 9.38% ± 1.14 to 6.55% ± 0.88 (1-tailed t-test, p = 0.036), and in layer 5 from 12.06% ± 1.29 to 7.82% ± 1.92 (1-tailed t-test, p = 0.045; Figure 2B, n = 8/group). There was a trend towards an overall decrease in TH+ axon density in all layers anterior to the injury site in the CCI-treated animals as compared to controls. TH+ axon density in the CCI tissue was reduced from 15.43% ± 2.94 to 10.04% ± 1.85 in layer 1, 8.88% ± 1.04 to 6.30% ± 1.15 in layer 2/3, and 14.48% ± 2.74 to 8.75% ± 2.55 in layer 5 (n = 8), respectively. However, these differences were not statistically significant (p = 0.07, 0.06, 0.07 in layers 1, 2/3, and 5, respectively; Figure 2A). Because many TH+ axons run from anterior to posterior in the the neocortex (Nakamura et al., 2000; Shnitko et al., 2014), it is expected that axons posterior to the injury site will be more strongly damaged than those anterior to the injury site. Indeed, this pattern of injury is similar to that we have previously reported using the identical CCI protocol and staining for serotonin axons using serotonin transporter immunoreactivity (Kajstura et al., 2018).

One month after injury, statistically significantly reductions are seen in TH+ axon density in all cortical layers both anterior and posterior to the injury site (Figure 3). Anterior to the injury site, where decreased TH+ axon density was not statistically significant at the 1 week time point, TH+ axon staining is now significantly reduced in the CCI animals as compared to the controls, decreasing from 17.33% ± 1.14 to 13.95% ± 0.79 in layer 1 (1-tailed t-test, p = 0.019), from 12.00% ± 0.63 to 8.55% ± 0.81 in layer 2/3 (1-tailed t-test, p = 0.0040), and from 10.60% ± 0.56 to 8.07% ± 0.84 in layer 5 (1-tailed t-test, p = 0.018; Figure 3A, n = 6/group). Posterior to the injury site, TH+ axon density remained significantly reduced in the CCI animals as compared to the control decreasing from 17.93% ± 1.30 to 10.05% ± 0.75 in layer 1 (1-tailed t-test, p = 0.00042), from 12.09% ± 0.52 to 7.86% ± 0.84 (1-tailed t-test, p = 0.0012), and from 11.26% ± 0.81 to 8.23% ± 0.69 in layer 5 (1-tailed t-test, p = 0.0091; Figure 3B, n = 6/group). Thus, by one month after injury, significant loss of TH+ axons density was found across neocortical layers and both anterior and posterior to the injury site.

**Figure 3.**
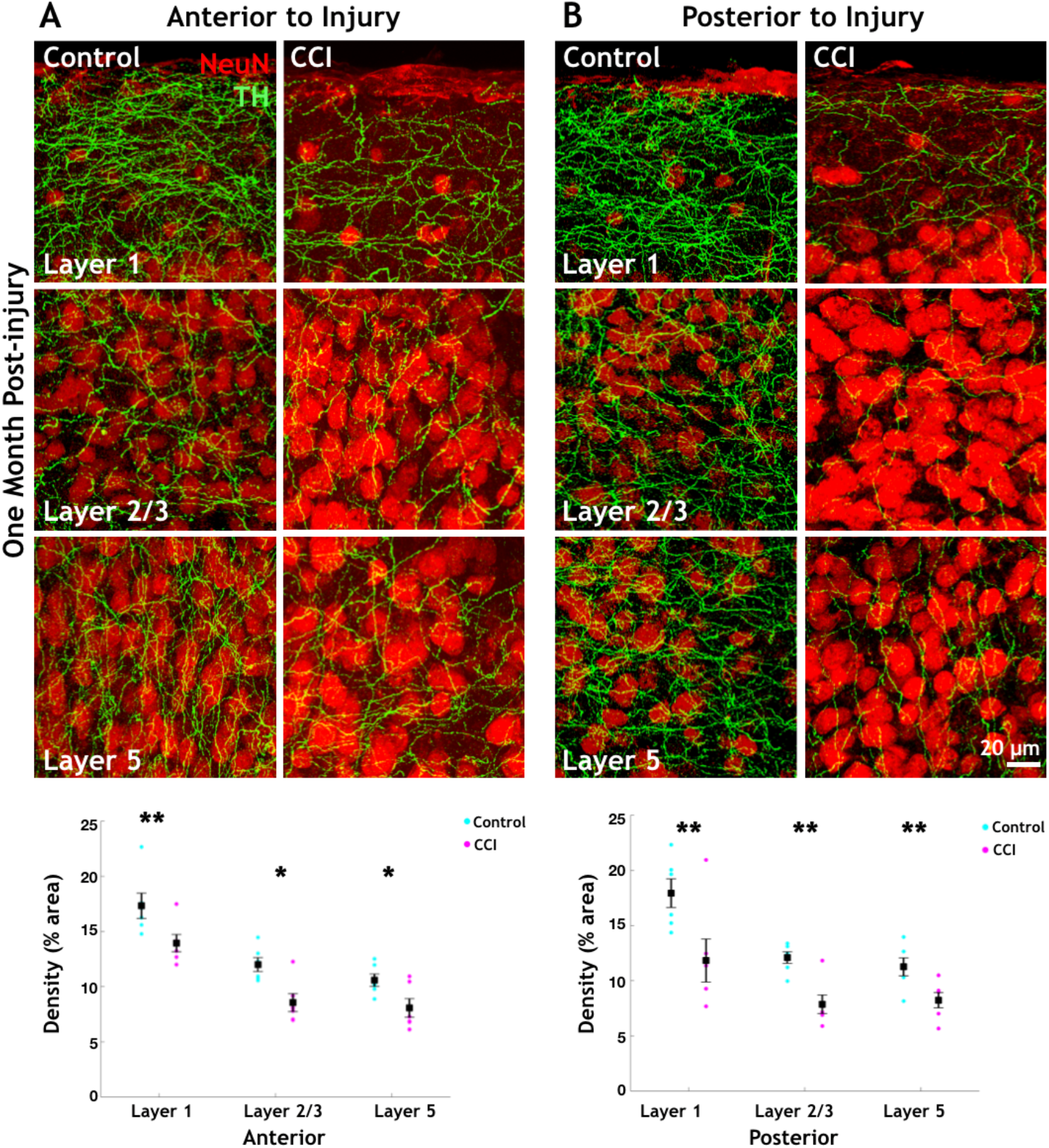
One month after CCI, the density of TH+ axons in the neocortex is reduced both anterior and posterior to the injury site to the injury site as compared with controls. **A.** Anterior to the CCI injury, significant loss of TH+ axon density is seen in cortical layers 1, 2/3, and 5. **B.** Posterior to the CCI injury, significant reductions in TH+ axon density are also observed in layer 1, layer 2/3, and layer 5. n= 6 animals/group. Onetailed Student’s t-test. * indicates p < 0.05. ** indicates p < 0.01. Error bars are SEM.

### Three months after CCI, TH+ axons in the neocortex regrew to recover their density to pre-injury levels

Three months after injury, significant differences in TH+ axon density between CCI and control animals had mostly been eliminated (Figure 4). Anterior to the impact injury, there were no statistically significant differences in TH+ axon density in any of the cortical layers surveyed. TH+ axon density in the CCI tissue was 21.53% ± 2.43 as compared to control 23.12% ± 2.50 in layer 1, 14.51% ± 1.72 and 17.07% ± 2.05 in layer 2/3 and, 21.31% ± 3.10 and 22.04% ± 3.04 in layer 5 (n = 8/group), respectively. These differences were not statistically significant (p = 0.34, 0.20, 0.44 in layers 1, 2/3, and 5, respectively; Figure 4A). Posterior to the impact injury, only cortical layer 2/3 remained at a significantly lower TH+ axon density as compared to control, down from 16.75% ± 1.91 in control tissue to 10.70% ± 1.15 in CCI (1-tailed t-test, p = 0.015; Figure 4B, n = 8/ group). TH+ axon density in the CCI tissue was 20.85% ± 3.16 as compared to control 28.51% ± 3.82 in layer 1, and 16.32% ± 1.78 and 20.76% ± 2.16 in layer 5 (p = 0.087 in layer 1, p = 0.082 in layer 5; Figure 4B)

**Figure 4.**
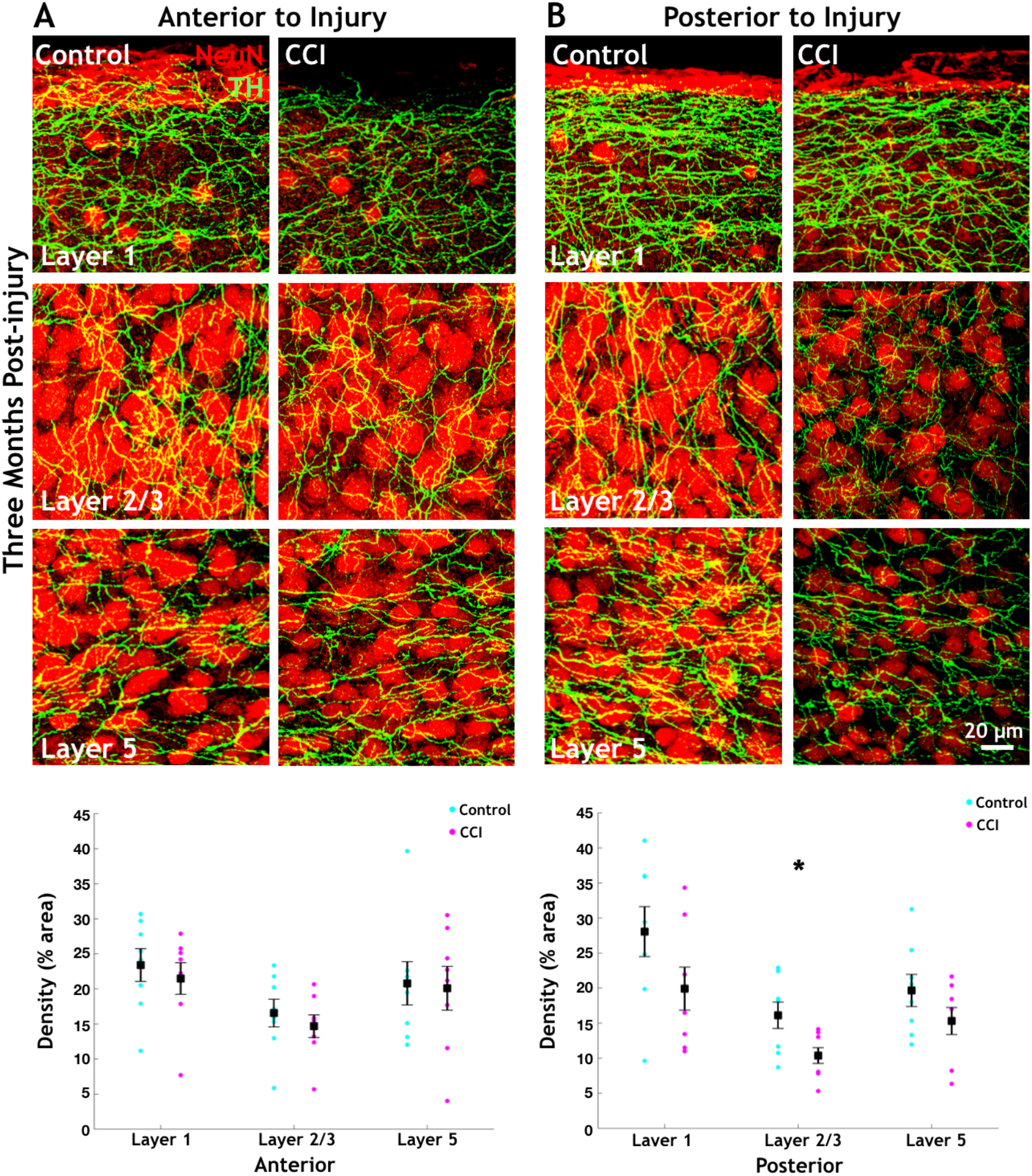
Three months after CCI, the density of TH+ axons in the neocortex is no longer significantly different from controls in most cortical layers. **A.** Anterior to injury TH+ axon density has normalized to the level of control tissue in all cortical layers examined. **B.** Posterior to injury significantly reduced TH staining is only seen in layer 2/3 of the CCI tissue. (n= 8 animals/group) One-tailed Student’s t-test. * indicates p < 0.05. Error bars are SEM.

Finally, we examined the time course of TH+ axon density changes, posterior to the CCI injury (Figure 5A, layer 1 shown). We compared density measurements from post-CCI time points (n = 6-8/group) to a pooled sample of control tissue from all time points (n = 22). In layer 1, we found a statistically significant loss of TH+ axons over time. TH+ axon density in the pooled control tissue group was 21.04% ± 1.89 in the CCI tissue density was reduced to 10.48% ± 1.38 at 1 day, 10.05% ± 0.82 at 1 month, and returned to 20.85% ± 3.38 at 3 months (ANOVA, F(3) = 5.9466, p = 0.0020; Figure 5B). We saw a significant reduction in TH+ axon density one week and one month post-injury as compared to the pooled control (1-tailed t-test, p = 0.0014 and 0.0052, respectively). Importantly, the TH+ axon density measured 3 months after CCI had recovered such that there was no significant difference between it and the pooled control group (1-tailed t-test, p = 0.48). In our view, this observation indicates that TH+ axons are capable of regrowth within 3 months of CCI injury. In layer 2/3, we found a statistically significant loss of TH+ axons over time (ANOVA, F(3) = 5.5825, p = 0.0028). In layer 5, we found a statistically significant loss of TH+ axons over time (ANOVA, F(3) = 5.6592, p = 0.0026).

**Figure 5.**
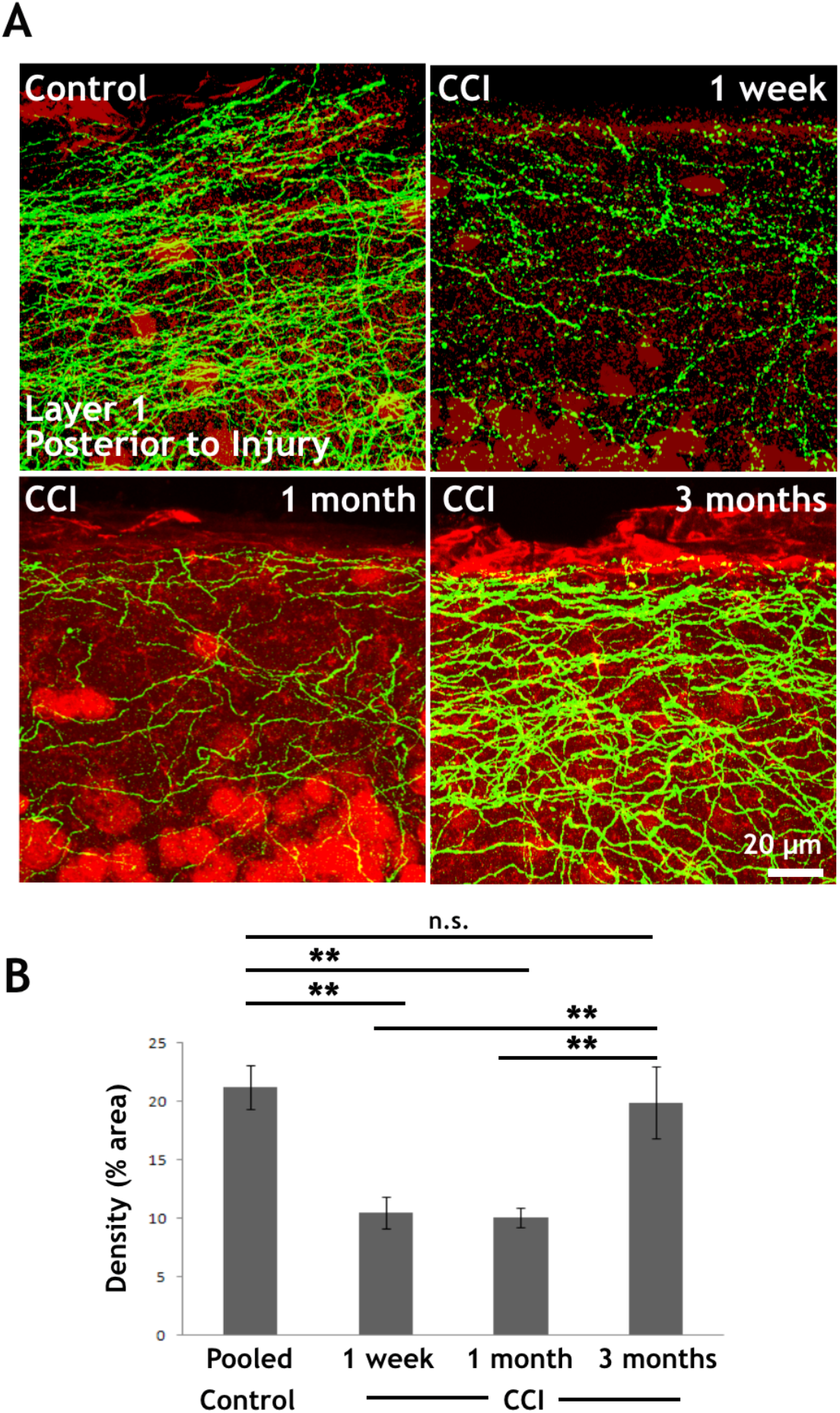
Images of neocortical layer 1, posterior to CCI injury, show loss and subsequent recovery of TH+ axon density. **A.** Exemplar images of TH+ axons in neocortical layer 1, posterior to injury, at three time points following CCI. Axon density is decreased at 1 week and 1 month after after injury but then returns to control levels by 3 months post-injury. **B.** Percent pixels within the image area positive for TH was compared in the CCI treatment over time (n= 6-8 animals/group) and also to a control sample set, pooled from all time points (n = 22). ANOVA followed by onetailed Student’s t-test ** indicates p< 0.01. Error bars are SEM.

### Subtypes of TH+ axons in the neocortex

As TH is present in both dopaminergic and noradrenergic axons in the neocortex (Weihe et al., 2006), it is useful to differentiate from which cell populations the axon damage and regrowth is occurring. In order to measure TH+ noradrenergic axons, we used dopamine-β-hydroxylase (DBH):cre BAC transgenic mice crossed with the mTmG reporter mouse line. We took this approach because we could not obtain antibodies directed against DBH that worked for immunohistochemistry. In this model DBH+ (presumed noradrenergic) neurons express membrane-targeted GFP (mGFP) and DBH-cells express membrane-bound Tomato (mTomato). We performed immunohistochemistry with the TH antibody in these mice using a far-red secondary and then quantified colabeling of the green (DBH-GFP) and far-red (TH) channels. In layer 1, of the somatosensory cortex 98.6% ± 0.8 of the TH+ axon segments were also DBH-GFP+ (n=149 axon segments from 3 mice). In layer 2/3, 96.6% ± 0.8 of the TH+ axon segments were also DBH-GFP+ (169 axons segments from 3 mice). In layer 5 however, only 78.9% ± 5.0 of the TH+ axon segments were also DBH-GFP+ (n= 326 axons segments from 3 mice). Similar patterns were seen in the motor cortex (not shown). These results suggest that the TH+ axons in the upper layers of the cortex are overwhelmingly noradrenergic while in the deeper layers both TH+ noradrenergic and TH+ non-noradrenergic axons are present (Figure 6A). We also performed immunohistochemistry with an antibody against the dopamine transporter (DAT) in wild type mice and saw moderate TH+ DAT+ axon density in the deep cortical layers while very sparse axonal staining is found in layers 1 and 2/3 (Figure 6B). This pattern of presumed dopaminergic and noradrenergic axon density is similar to that previously reported in the neocortex of rats and mice (Berger et al., 1976; Levitt and Moore, 1978; Papadopoulos and Parnavelas, 1991; Berger et al., 1991; reviewed in Schubert et al., 2015).

**Figure 6.**
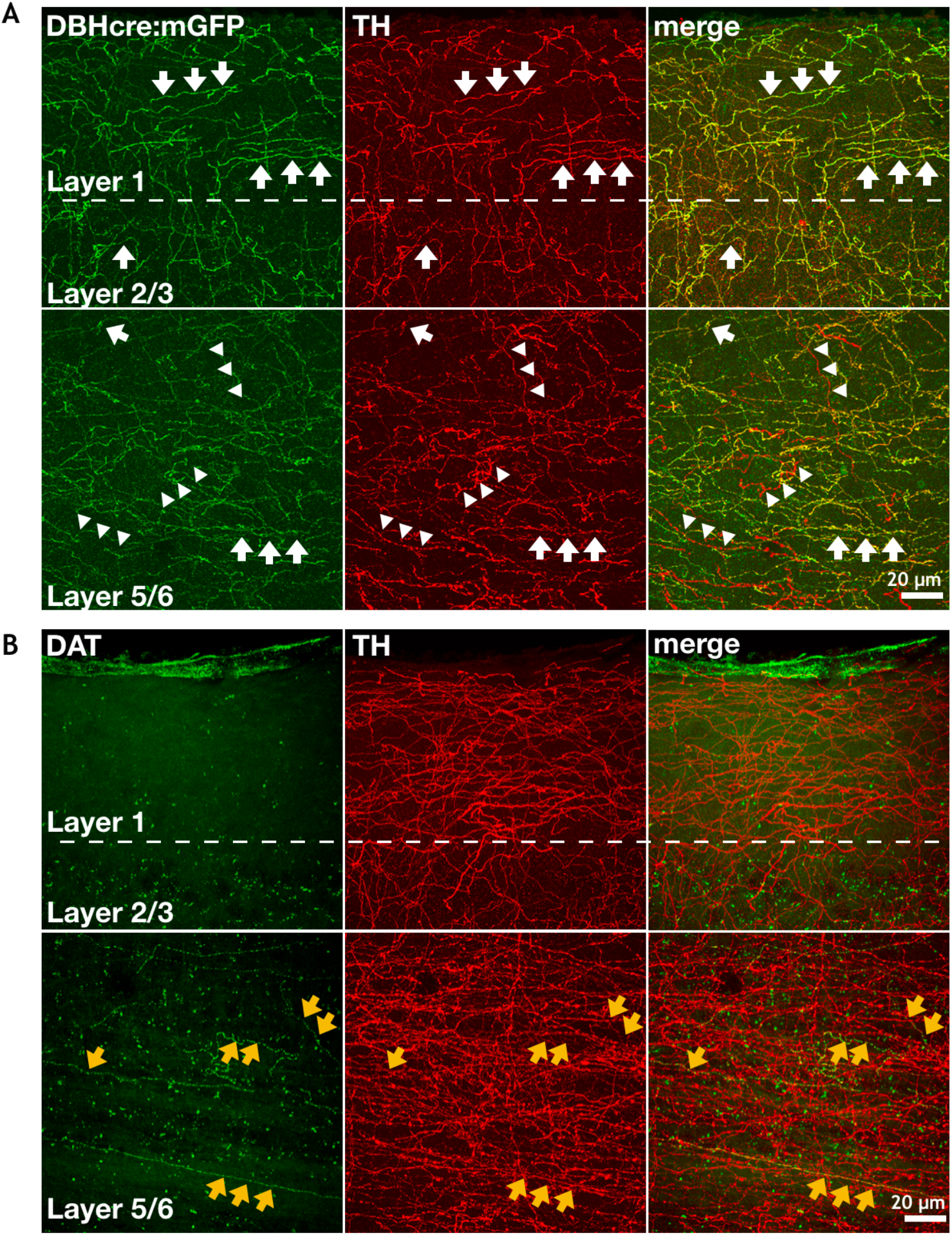
Double label immunohistochemistry shows that TH+ axons in the neocortex are also immunopositive for dopamine beta hydrolase-GFP (DBH-GFP) or the dopamine transporter in uninjured tissue. **A.** Exemplar images with DBH-GFP expression in green and TH immunoreactivity in pseudocolored red. In the superficial cortical layers (L1 and L2/3) nearly all TH+ axons are also DBH-GFP+ (white arrow show examples). However, in layer 5, in addition to TH+/DBH-GFP+ axons (white arrows), TH+/DBH-GFP-axons are present as well. (white arrowheads). **B.** Exemplar images with dopamine transporter (DAT) immunoreactivity in green and TH immunoreactivity in pseudocolored red. In the superficial cortical layers (L1 and L2/3) DAT+ staining is quite rare while in the deep cortical layers some DAT+ axons are clearly evident. (some TH+/DAT+ axons are indicated with yellow arrows)

### VAChT+ axons in the neocortex were not damaged by CCI

Cholinergic fibers were visualized with an antibody directed against the vesicular acetylcholine transporter (VAChT) which is found within the axons of cholinergic neurons (Ichikawa et al., 1997). Importantly the overall trajectory of cholinergic axons innervating the neocortex is different from that of monoamine axons (Figure 1A; Paul et al., 2015) as the former tend to ascend in the ventral-to-dorsal orientation.

VAChT axon density was unaltered by CCI treatment, both anterior and posterior to the injury site at one week post-injury, in all cortical layers measured (n = 8/group, Figure 7). These tissue slices were from the same mice used to assess TH+ immunoreactivity. There were no statistically significant differences in the CCI group as compared to control following injury (1-tailed t-tests anterior: p = 0.42, 0.27, 0.16 in layers 1, 2/3, and 5, respectively, Figure 7A; posterior: p = 0.17, 0.14, 0.21 in layers 1, 2/3, and 5, respectively, Figure 7B). Although loss of TH+ axons was pronounced by the one month post-injury time point (Figure 3), VAChT axons remained spared. There were no statistically significant differences in the CCI group as compared to control one month or three months following injury (one month anterior: p = 0.46, 0.34, 0.50 in layers 1, 2/3, and 5, respectively, posterior: p = 0.46, 0.38, 0.32 in layers 1, 2/3, and 5, respectively (one month, n = 6/group); three months anterior: p = 0.30, 0.31, 0.25 in layers 1, 2/3, and 5, respectively, posterior: p = 0.45, 0.27, 0.44 in layers 1, 2/3, and 5, respectively (three months, n = 8/group); Supplemental Figure 1). The VAChT immunohistochemistry images herein, and relative staining pattern of cortical cholinergic fibers, are consistent with previous reports in the neocortex of rats and mice (Arvidsson, 1997; Bloem et al., 2014).

**Figure 7.**
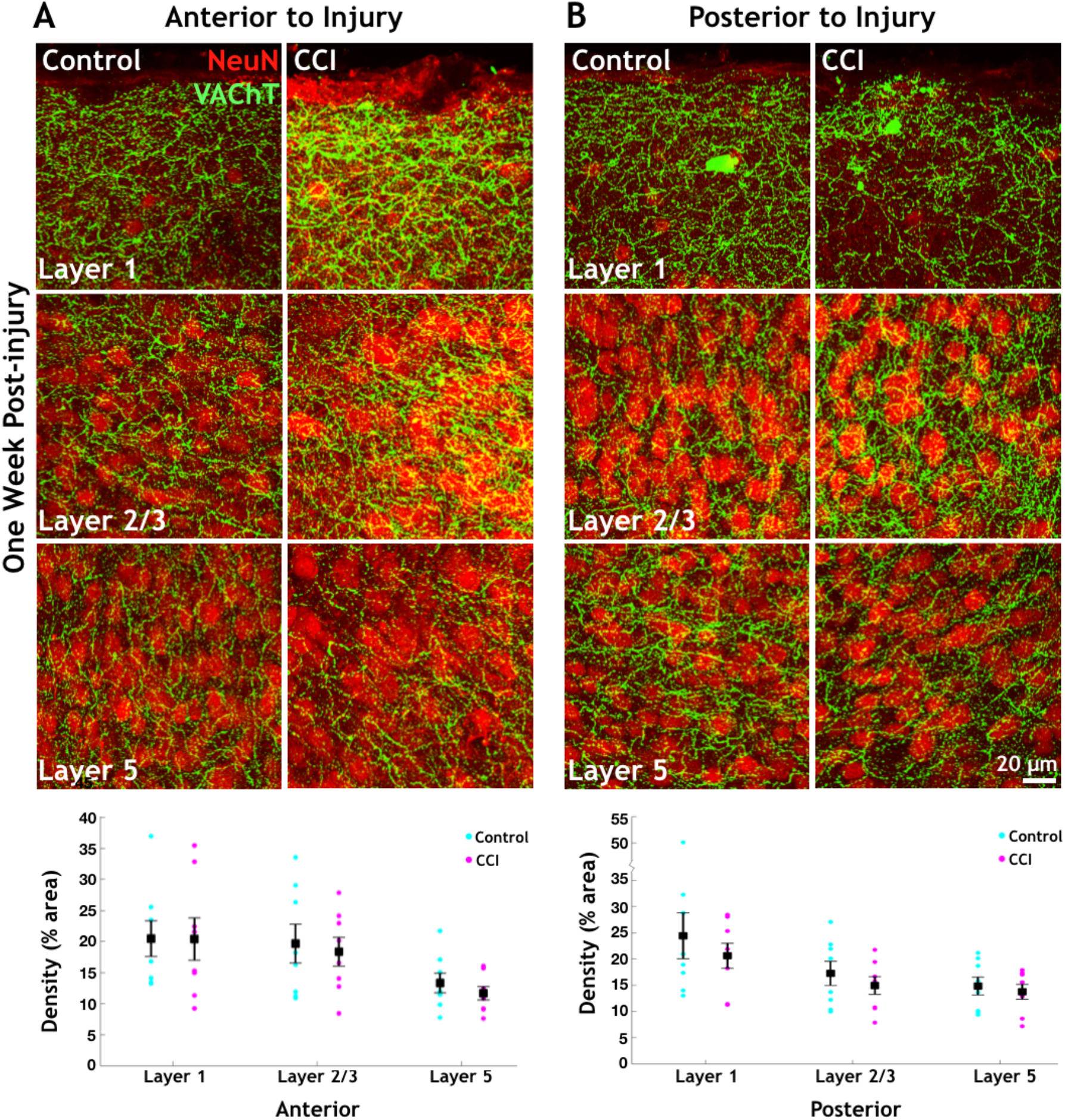
One week after CCI, the density of VAChT+ axons in the neocortex is not significantly changed when compared with controls. **A.** Maximally projected 40x confocal z-stacks with thresholded VAChT staining for acetylcholine axon labeling and NeuN staining for cell bodies to indicate cortical column location. Anterior to injury no overt loss of VAChT+ axons is observed in any cortical layer examined as compared to control tissue. **B.** Posterior to injury no overt loss of VAChT+ axons is observed in any cortical layer examined as compared to control. No statistically significant changes in VAChT+ axon density are observed either anterior or posterior to the injury site. (n= 8 animals/group) One-tailed Student’s t-test. Error bars are SEM.

### TH+ axons in the neocortex were damaged by a stab injury, but showed regrowth when measured 1 month afterwards

To assess whether TH+ axons were capable of regrowth following an additional form of injury, we performed axon-transecting cortical stabs through all layers of the somatosensory cortex of adult mice. We used a blade oriented in the coronal plane to preferentially interrupt axons running in an anterior-to-posterior direction. We then performed immunohistochemistry to reveal TH+ axons at various times after the stab injury. Low magnification images were first acquired to assess the overall extent of the stab injury and its recovery over time (Figure 8A-C). Fixed-tissue stab rift size varies as a result of slice processing but approximate areas of injury were identified via NeuN-labeled cellular debris halos (not shown) and TH+ axon loss (Figure 8A-C, dotted yellow lines). Single-photon confocal 35 μm thick z-stacks were then acquired using a 40x objective in layers 2/3 and 5 of the somatosensory cortex including the stab rift. Maximum-projected images were analyzed for TH+ axon density. TH+ stained fibers within the central 50 μm portion of the rift were exhaustively traced (Figure 8D-I, solid blue lines) and compared across injured and uninjured contralateral tissue. Total axon length within the central rift zone was quantified. In cortical layer 2/3, following stab-induced TBI, axon length within rift averaged 636.5 μm ± 309.2, 819.2 μm ± 158.6, and 1404.4 μm ± 191.1, at one day, one week, and one month, respectively. In the uninjured contralateral hemisphere, axon length within the corresponding tissue volume was 1632.6 μm ± 110.5. There was a significant increase in TH+ axon density in layer 2/3 over time following stab injury (ANOVA, F(3) = 6.9033, p = 0.0104). Posthoc student’s t-tests revealed significant differences between the average TH+ axon length between layer 2/3 at one day and one month (p = 0.040) and between one week and one month (0.028). Axonal length traced was not different between one day and one week (p = 0.31). Statistically significant differences were also observed between the stabbed and contralateral control hemisphere TH+ axon density at one day (p = 0.012), and at one week after injury (p = 0.0028). Traced axon length was not significantly different between one month and the contralateral control region (p = 0.17; n = 4/group).

**Figure 8.**
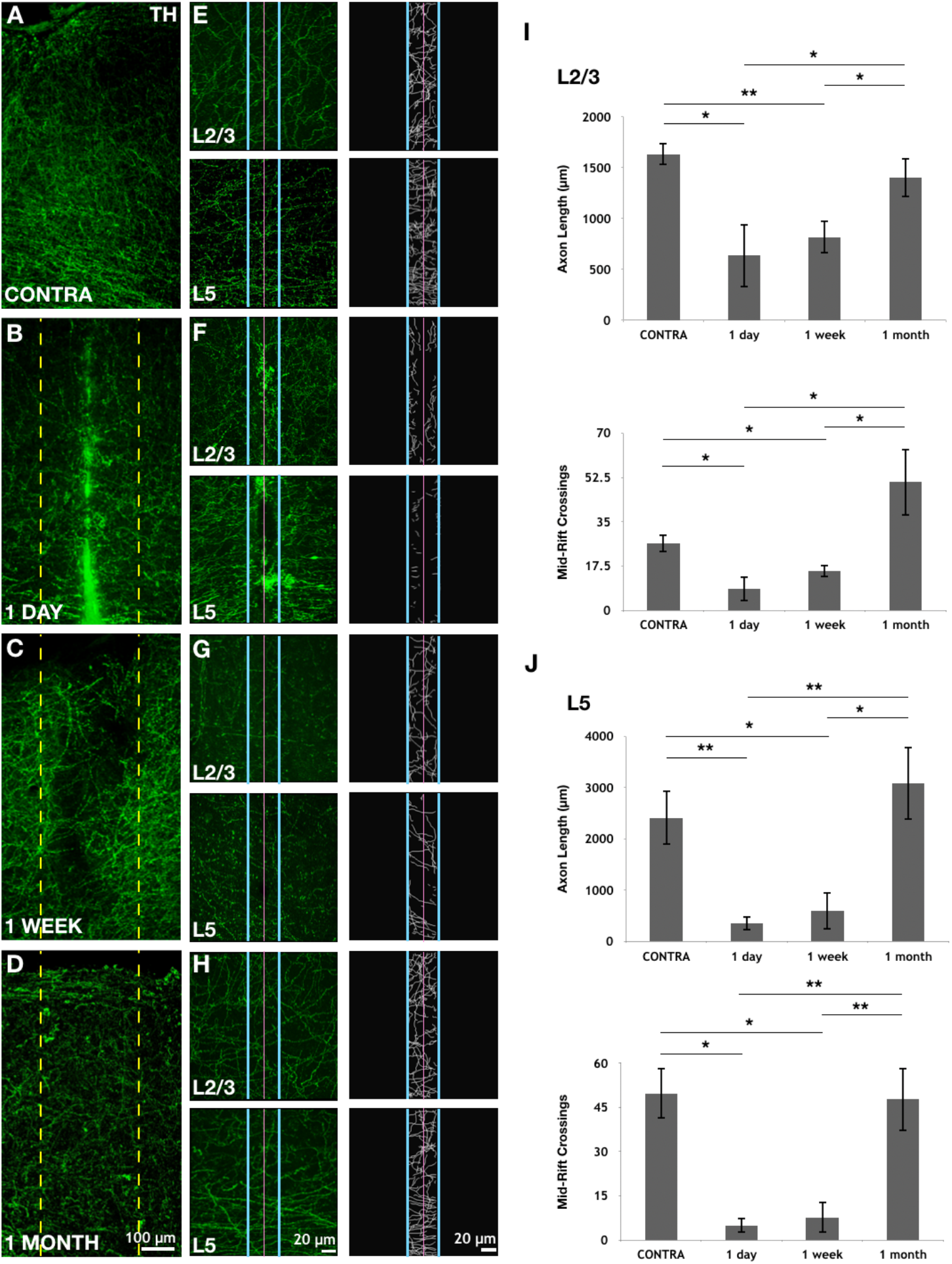
TH+ axons are damaged by a neocortical stab injury but recover within 1 month. Maximally projected confocal z-stacks from the somatosensory cortex with TH staining for catecholaminergic axon labeling in pseudocolored green. **A.** 10x z-stack exemplar image of the contralateral, uninjured hemisphere. **B-D.** Exemplar images one day, one week and one month and after injury. The yellow dotted lines indicate the approximate stab rift region. **E.** The left column shows TH+ immunoreactive axons in the contralateral, uninjured hemisphere in layers 2/3 and 5. The right column shows exhaustive axon tracing within a region of interest bounded by blue vertical lines. The pink vertical line is the midline of this region of interest. **F-H.** Exemplar images and tracing, using the organizing scheme of panel E, for time points one day, one week and one month after tab injury. **I.** Quantification of exhaustively traced axon length and center rift (pink line) crossings in layer 2/3. **J.** Quantification of exhaustively traced axon length and center rift crossings in layer 5. (n = 4 animals/group) ANOVA followed by one-tailed Student’s t-test * indicates p< 0.05 and ** indicates p < 0.01. Error bars are SEM.

In cortical layer 5, following stab injury, axon length within rift averaged 353.6 μm ± 135.0, 592.4 μm ± 367.4, and 3080.6 μm ± 710.9, at one day, one week, and one month, respectively. In the contralateral hemisphere axon length within the same tissue volume was 2412.2 μm ± 533.3. There was a significant increase in TH+ axon density over time in layer 5 following stab injury (ANOVA, F(3) = 7.8773, p = 0.0069). Posthoc student’s t-tests revealed significant differences between the average TH+ axon length in layer 5 at one day and one month (p = 0.0047) and between one week and one month (p = 0.010). Axonal length traced was not difference between one day and one week (p = 0.28). Statistically significant differences were also observed between one day and the contralateral hemisphere TH+ axon density (p = 0.0048), and one week and contralateral (p = 0.015). Traced axon length was not significantly different between one month and contralateral (p = 0.24; n = 4/group). Layer 1 was not analyzed because it was not completely intact in every stab sample.

We also analyzed the number of axons crossing the center of the rift in each image. In cortical layer 2/3, following injury, the number of axons crossing the rift averaged 8.5 ± 4.8, 15.5 ± 2.4, and 50.75 ±13.3, at one day, one week, and one month, respectively. In the contralateral hemisphere mid-rift crossings through the center of the same tissue volume was 26.5 ± 3.6. In cortical layer 5, following stab-induced TBI, traced-filaments crossing the rift averaged 5.0 ± 2.6, 7.7 ± 5.2, and 47.7 ± 10.7, at one day, one week, and one month, respectively. In the contralateral hemisphere mid-rift crossings through the center of the same tissue volume was 49.7 ± 8.6. There was a significant increase in TH+ axons crossing the rift over time following stab injury in layer 2/3 (ANOVA, F(3) = 5.7908, p = 0.0174) and layer 5 (ANOVA, F(3) = 8.9509, p = 0.0046). Posthoc student’s t-tests revealed significant differences between the average TH+ axon crossings in layer 2/3 at one day and one month (p = 0.021), between one week and one month (0.037), between one day and contralateral (p = 0.014), and between one week and contralateral (p = 0.024). Axonal crossings were not significantly different between one day and one week (p = 0.13) nor did they differ between one month and contralateral (p = 0.08). Statistically significant differences were also present between the average TH+ axon crossings in layer 5 at one day and one month (p = 0.012), between one week and one month (p = 0.013) between one day and contralateral (p = 0.0051), and between one week and contralateral (p = 0.0047). Axonal crossings were not different between one day and one week (p = 0.31) nor did they differ between one month and contralateral (p = 0.44). These results indicate that, following stab injury, TH+ axons are able to regrow in both layers 2/3 and 5, at least in part by passing across the stab rift. Serotonin transporter-mGFP+ axons were also observed in these slices and showed stab-induced damage and subsequent regrowth (not shown) as previously reported (Jin et al., 2016).

## Discussion

The main finding of this investigation is that two different forms of trauma delivered to the neocortex of adult mice, CCI and stab injury, produced a reduction in the density of catecholaminergic axons, measured using TH immunohistochemistry, followed by a slow recovery on the time scale of weeks-to-months.

One week after CCI, the density of TH+ axons was significantly reduced across the neocortical layers in the tissue lying posterior to the injury as compared to controls (Figure 2). By one month after injury, significant reductions in TH+ density were seen across cortical layers both anterior and posterior to the injury site (Figure 3) suggesting a delayed loss of axons in the anterior region produced by retrograde degeneration, as the TH+ axons largely run in an anterior-to-posterior direction in the neocortex (Figure 1A; Nakamura et al., 2000; Shnitko et al., 2014). Importantly, by 3 months after CCI injury, significant differences in TH+ axon density between injured and control tissue were no longer present, save in layer 2/3 of the posterior compartment (Figures 4 and 5). This pattern of loss and recovery of TH+ axons density is similar, but not identical to that previously observed using measurements of serotonin axon density (revealed by serotonin transporter immunohistochemistry) together with the identical CCI and control treatments: Serotonin axon density was significantly reduced 1 week and 1 month after CCI only in the posterior compartment and recovery of serotonin axon density was well underway at the 3 month time point, with no significant differences remaining between CCI and control tissue (Kajstura et al., 2018).

Might the apparent loss and regrowth of TH+ axons following CCI, indicated by TH+ immunohistochemistry, be an artifact produced by nonspecific tissue damage? The observation that, at the one month postinjury time point, the tissue anterior to the CCI injury showed significantly reduced TH+ axon density (Figure 3) but not serotonin axon density (Kajstura et al., 2018), argues against a non-specific reduction in axonal immunopositive signals. Likewise, the failure of CCI injury to produce significant reductions in cholinergic axon density, indicated by VAChT immunohistochemistry, in any neocortical layer, either anterior or posterior to the CCI injury site (Figure 7 and Supplementary Figure 1) argues that the loss and recovery of TH+ axons seen here following CCI was a specific effect.

When TH+ axons were transected by a stab injury, both the total length of axons and the number of axons crossing the center of the rift were significantly reduced as compared with the uninjured contralateral side. These reductions were apparent in both layer 2/3 and layer 5 and were similar when measured 1 day or 1 week after the stab injury. However, when measurements were made 1 month after injury, both the total length within the rift zone and the number of center rift crossings of TH+ axons had returned to or exceeded control (uninjured contralateral side) levels (Figure 8). These results, transient axon disruption with minimal retrograde degeneration, followed by regrowth within 1-2 months is similar to those previously seen when serotonin axon immunohistochemistry or live imaging was performed using the same stab protocol (Jin et al., 2016).

Taken together, these findings suggest that catecholaminergic axons in the neocortex of the adult mouse can regrow following two different glia scar-forming injuries. There are, however, some important caveats to be sounded in relation to these findings. First, TH is present in both dopamine and norepinephrine-using neurons (Weihe et al., 2006) and so TH+ axons may potentially be either noradrenergic or dopaminergic. Our double staining results in DBHcre:mGFP norepinephrine marker mice show that ~97% of TH+ axon segments in layers 1-3 of the somatosensory neocortex are also DBH-GFP+. Furthermore, only ~2% of TH+ axon segments in layers 1-3 in WT mice were also immunoreactive for the dopamine transporter. In layers 5-6, however, ~79% of the TH+ axon segments were also DBH-GFP+ and ~9% of the TH+ axon segments were DAT+ (Figure 6). In our view, the large loss and regrowth of TH+ axons in layers 1-3, where ~97% of them are DBH-GFP+, suggests that noradrenergic axons are damaged and regrow following both CCI and stab injuries. The loss and regrowth of axons in layer 5 is harder to interpret as there are notable populations of both DBH-GFP+ and DAT+ axons in this layer. The loss and regrowth of TH+ axons in layer 5 could result from changes in noradrenergic axons alone, or both noradrenergic and dopaminergic axons alone. Thus, whether dopaminergic axons are damaged and regrow following CCI or stab injuries remains unclear.

A second caveat is that histological experiments are snapshots and so are unable to provide us with dynamic information. Is the recovery of TH+ axon density following injury due to growth from the damaged ends of axons (regeneration) or is it due to local sprouting (new growth originating from undamaged axons; definitions from Tuszynski and Steward, 2012)? When TH+ axons regrow, do they follow the pathways abandoned by TH+ axons that previously degenerated? Is the ultimate survival time of regrown axons comparable to surviving or uninjured axons? Are there morphological signatures that predict which TH+ axons will survive CCI injury? Answering these questions will require long-term *in vivo* imaging experiments.

A third caveat is that we don’t know whether the regrown TH+ axons function normally. Do they convey action potentials and if so, do they do so with the same fidelity? Are the axonal varicosities of the regrown axons competent to release neurotransmitter and do they do so with similar probability and short-term dynamics in response to action potentials? Newly developed optical probes for norepinephrine (Feng et al., 2019) and dopamine release (Patriarchi et al., 2018; Sun et al., 2018) hold promise to address these questions when combined with *in vivo* TH+ axon imaging.

Glutamatergic axons in the adult mammalian CNS have a very limited capacity to regrow after injury (Heubener and Strittmatter, 2009; Canty et al., 2013; Akassoglou et al., 2017). Glutamate, along with GABA and glycine are neurotransmitters that activate ionotropic receptors at classical synapses in which the presynaptic active zone faces a receptor-laden postsynaptic density lying across the narrow synaptic cleft. This form of point-to-point neurotransmission is built to convey information quickly (on a time scale of milliseconds to tens of milliseconds) and in a very spatially restricted way (with transmitter spillover to only a few adjacent synapses). The monoamine neurotransmitters serotonin, norepinephrine and dopamine mostly act via metabotropic receptors. As a consequence, they convey information on a slower timescale, appropriate for conveying modulatory information about internal states like mood, appetite or libido. Monoamine releasing axonal varicosities can sometimes form conventional point-to-point synapses, but their action in the neocortex is dominated by volume transmission (Seguela et al., 1990; Umbriaco et al., 1995; Fuxe et al., 2007; Decarries et al., 2008). In volume transmission, there is no postsynaptic density bearing a high concentration of receptors facing the presynaptic active zone. As such, monoamine neurotransmitters must diffuse in the extracellular space for a much greater distance to encounter their dispersed receptors.

As a consequence of point-to-point transmission, if a damaged glutamate axon regrows, it must reinnervate its postsynaptic targets with great accuracy in order to reconstitute pre-injury function. By contrast, volume transmitting axons that regrow need only target the general area that was denervated, as their signaling is not so spatially or temporally constrained. Perhaps, volume transmitting axons, like those of the monamine transmitters, have been endowed with the capacity to regrow after injury to the adult brain because such regrowth would restore pre-lesion function even if it does not recapitulate the precise synaptic targeting of the previously injured axon.

What are the molecular specializations that allow serotonin and catecholamine axons to regrow while other injured axons in the brain fail to do so? Gene expression profiling of regrowth-competent neurons before and following injury should be informative in this regard, as it has been in the peripheral nervous system (Li et al., 2015; Dulin et al., 2015; He and Jin, 2016; Pan et al., 2017). A useful strategy for uncovering the molecular mechanisms underlying axon regrowth in the brain may to be search for those basal and injury-evoked gene expression events that are shared by monoaminergic neurons. By uncovering the unique molecular properties of regrowth-competent neurons in the brain, we hope to inform the development of therapies to promote axon regrowth in a wide variety of cell types and thereby aid functional recovery following brain injury.

## Acknowledgments

These experiments were inspired by the long-standing investigations of our colleague Mark Molliver, who died in 2012. Thanks to members of the Linden lab, Jeremiah Cohen, and Michele Pucak for useful suggestions. Funding was from the Johns Hopkins University Brain Science Institute and NIH NS81467 (D.J.L.), and NIH F32 NS090822 (S.E.D.). The Johns Hopkins Imaging Core Facility, which aided this effort, is supported by NIH NS050274.

## Author contributions

S.E.D. designed experiments, performed experiments, analyzed data, and wrote the paper. T.J.K. designed and performed experiments. Y.J. designed experiments and provided intellectual contribution. M.H.C-C. performed imaging experiments and analyzed data. A.K. performed experiments. D.J.L. designed experiments and wrote the paper.

---

**Supplemental Figure 1.**
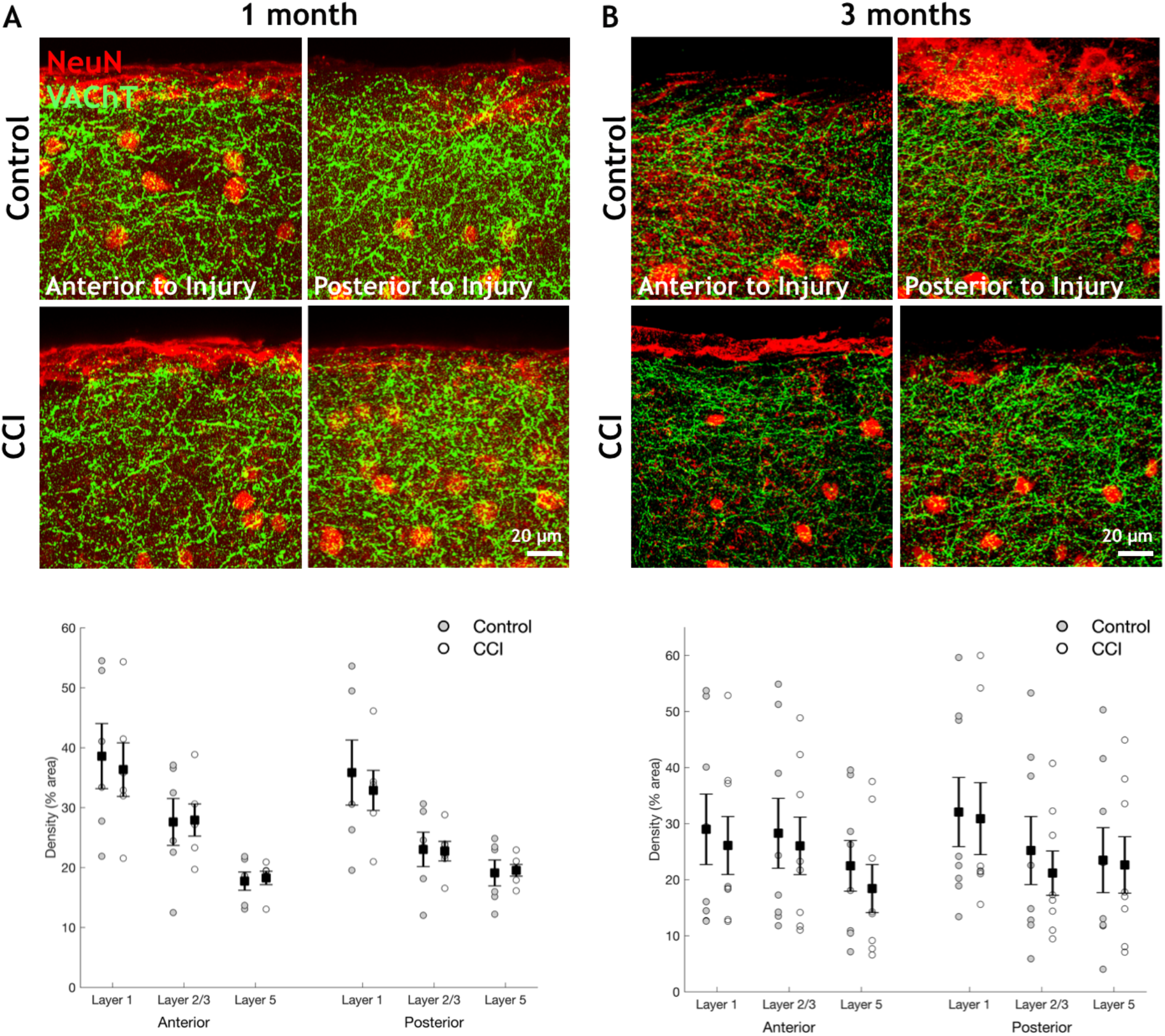
One month or three months after CCI, the density of VAChT+ axons in the neocortex is not significantly changed when compared with controls. **A.** Exemplar images are from are cortical layer 1. At one month post-injury, no statistically significant changes in VAChT pixel density are observed, either anterior or posterior to the injury site. **B.** Exemplar images are from are cortical layer 1. At three months postinjury, no statistically significant changes in VAChT pixel density are observed, either anterior or posterior to the injury site. (n= 6-8 animals/group) One-tailed Student’s t-test * indicates p < 0.05 and ** indicates p < 0.01. Error bars are SEM.

